# Sex determination system turnovers play important roles in the willows speciation

**DOI:** 10.1101/2023.10.23.563523

**Authors:** Zhi-Qing Xue, Wendy L. Applequist, Elvira Hörandl, Li He

**Affiliations:** Eastern China Conservation Centre for Wild Endangered Plant Resources, Shanghai Chenshan Botanical Garden, Shanghai, China; William L. Brown Center, Missouri Botanical Garden, St. Louis, Missouri 63110, U.S.A; Department of Systematics, Biodiversity and Evolution of Plants (with Herbarium), University of Goettingen, Göttingen, Germany

**Keywords:** *Salix*, Sex determination, Speciation, Sex chromosomes turnover, Genome evolution

## Abstract

Almost all species in the genus *Salix* (willow) are dioecious, but some have male and some female heterogamety, and the chromosomal location of the sex-linked regions (termed SDSs) differs between different species. We first analyzed the SDSs of two species, *Salix cardiophylla* and *S. interior*, whose positions in the *Salix* phylogeny make them important species for understanding a sex chromosome turnover that has been detected in their relatives, and that changed the system from male to female heterogamety. We show that both species have male heterogamety, with XY-linked regions on chromosome 15 (termed a 15XY system). The sex-linked regions occupy 21.3% and 22.8% of the entire reference chromosome, respectively. By constructing phylogenetic trees of species with known SDSs, we determined the phylogenetic positions of all the species. Reconstruction of SDSs revealed that 15XY system is likely the ancestral of willows. Finally, we tested for both current and ancestral gene flow between different species with the same or different sex-determining systems, as the sex chromosomes can play important roles in reproductive isolation between species. We inferred lower gene flow between species with XY on chromosome 7 (7XY) and ZW on chromosome 15 (15ZW) systems, compared with gene flow either between species with XY on chromosome 15 (15XY) and 15ZW systems or between species with 7XY and 15XY systems. We argue that, although sex chromosomes turnovers in willows may not create complete reproductive barriers, gene flow may be reduced between species with different SDSs.

## Background

Chromosome rearrangements can suppress recombination and may enhance reproductive isolation between populations with different arrangements during speciation, and between genes contributing to isolation through adaptive mutations that interact to create Bateson–Dobzhansky-Muller incompatibilities (Baack et al., 2015). Moreover, compared with the autosome, sex chromosomes can form barrier to introgression between diverging lineages and thus play a special role in speciation, which mainly due to four main factors (Qvarnström & Bailey, 2009): (1) relative speed of evolution in ecological traits differed on sex chromosomes versus autosomes, while this may only involve relatively few sex-linked genes; (2) non-random accumulation of genes coding for the sex-specific fitness traits on sex chromosomes; (3) exposure of incompatible recessive genes in hybrids, for example, Graves (2016) thought that when autosomes in one species have become sex chromosomes in another, hybrids between individuals with different sex chromosome pairs are likely to suffer from infertility because of pairing and recombination disruptions or the heterospecific interactions between genes that determine fertility on the sex chromosomes; (4) lower recombination in heterogametic sex chromosomes. Sex chromosomes are also involved in Haldane’s Rule in animals that the heterogametic sex (XY or ZW) is more likely to be affected in an interspecific cross, and large X effects that the disproportionately large role of the X chromosome in reducing hybrid fitness have been detected in many taxa (Faria & Navarro, 2010; Graves & O’Neill, 1997; Presgraves, 2018). Sex determination systems can involve male heterogamety (often termed XX/XY systems, even when there is no completely sex-linked region, or only a small one) or female heterogamety (ZW/ZZ). Sex chromosomes in plants have evolved in many lineages, and can differ between close-related species (Slancarova et al., 2013; Tennessen et al., 2018), similar to the young sex chromosome systems known in some animal groups, including amphibia and fish (Dufresnes et al., 2020; Franchini et al., 2018). In angiosperms, shifts between the XY and ZW systems are known in several genera, including *Dioscorea*, *Silene*, *Populus*, and *Salix* (Leite Montalvão et al., 2021). Such changes, and the changes of chromosomal location, known as turnover events, could be important in speciation (Tennessen et al., 2018). Sex chromosome turnovers are termed homologous when same sex chromosome pair, remains unchanged (as in the change from male to female heterogamety on the chromosome 19 in the genus *Populus* (Salicaceae; Hyden et al., 2021). “Nonhomologous transition” (or “trans-heterogamety transitions”) refer to changes in which a sex-determining gene emerges on an autosome (Kondo et al., 2006), including transposition of an existing male-determination locus to an autosome (Yang et al., 2020). In the *Salix*, two different chromosomes (7 and 15) are known to carry sex-determining loci (Wang et al., 2023a). In addition, chromosomal fusions can also cause sex chromosome turnover, as in the plant *Rumex hastatulus*; in this species, an X-autosome fusion created a new sex chromosome morphology (and probably affected recombination on the chromosomes). It has been proposed that this contributed to reproductive isolation, since the neo-sex chromosome origin is inferred to have coincided temporally with cessation of gene flow in the two races whose populations have the different sex chromosome cytotypes (Beaudry et al., 2020). Such trans-heterogamety transitions might promote speciation especially if genes that control reproductive isolation between populations accumulate on neo-sex chromosomes, as they are though to do (see above) on the ancestral sex chromosomes (Kitano & Peichel, 2012).

These arguments suggest that turnovers could lead to accelerated speciation. Hybridization and admixture among lineages with different SDSs have rarely been studied in plants while this has been widely studied in animals (Dixon et al., 2019; Kuwana et al., 2021; Miura et al., 2012; Natri et al., 2019). Species of toads (Bufonidae) with different sex-determining systems (*Bufo bufo* with ZW system and *B. spinosus* with XY system) could hybridize only across a narrow area (10km wide) where reproductive isolation (genetic incompatibilities) likely prevents their gene pools from merging (Dufresnes et al., 2020). Both XY and ZW sex determination systems (on multiple different chromosomes) were found in East African cichlids, and rapid turnovers could be associated with the rapid adaptive radiations in different lakes (Feller et al., 2021). Plant species of the section *Otites* in genus *Silene* with both XY (*S. colpophylla* and *S. latifolia*) and ZW systems (*S. otites* and *S. borysthenica*), the change in heterogamety of this section might contribute from a species with the male-determining chromosome hybridize with another species (Balounova et al., 2019). The effects of sex chromosome turnover events in phenotypic diversification and speciation are not yet clear (El Taher et al., 2021).

Taxonomic groups with diverse sex chromosomes, with both male (XY) and female (ZW) heterogamety, can help understanding whether, and how, sex-linked regions may affect reproductive isolation and speciation (Filatov, 2018; Ogata et al., 2021). The willow genus, *Salix* (Salicaceae, which is closely related to *Populus*) includes two major clades, *Salix* and *Vetrix*. Species of the *Salix* clade usually have XY SDS on chromosome 7, while those of clade *Vetrix* have both XY and ZW SDSs on chromosome 15. The SLR has therefore shifted between chromosomes as well changes heterogamy (Gulyaev et al., 2022; Wang et al., 2023a), making *Salix* ideal for studying the relationship between sex chromosome turnovers and speciation. In particular, in the *Salix* clade, *Salix nigra, S. chaenomeloides,* and *S. dunnii* have XY SDSs on chromosome 7, henceforth abbreviated to 7XY (He et al., 2021; Sanderson et al., 2021; Wang et al., 2022a). Most studied species of the *Vetrix* clade, *S. suchowensis* (Hou et al., 2015), *S. viminalis* (Almeida et al., 2020), *S. purpurea* (Zhou et al., 2020), *S. polyclona* (He et al., 2023a), *S. koriyanagi*, *S. integra*, and *S. udensis* (Wilkerson et al., 2022), have ZW SDSs on chromosome 15 (15ZW systems). However, three basal species in the *Vetrix* clade (*S. arbutifolia*, *S. triandra*, and *S. exigua*) have 15XY systems (Hu et al., 2022; Wang et al., 2022a; Wang et al., 2023a). The SDSs of other basal species, including *S. cardiophylla* and *S. interior*, are still unknown (Gulyaev et al., 2022; Wu et al., 2015).

Hybridization events in *Salix* are also common (Percy et al., 2014). Natural hybridization is usually between species in the same clade (Wagner et al., 2021). However, several species can produce seeds in crossing experiments. For example, *S. cardiophylla* and *S. arbutifolia* (15XY) in the *Vetrix* clade can produce fertile offspring, which are named *S. kamikotica* (Kimura, 1931, 1937). In addition, S*alix exigua,* a diploid species (15XY) showed no pollination barrier in crosses with other diploid representatives of the same *Vetrix* clade (*S. eriocephala*, and *S. petiolaris*), and produced fertile *F*_1_ offspring (Boufford, 1993; Mosseler, 1990). Fertile crosses have also been recorded between *S. triandra* (15XY) and *S. viminalis* (15ZW) and produced the F1 hybrids of *S. mollissima* (Gulyaev et al., 2022; Karp et al., 2011), while the hybrids were equally susceptible compared with the parents (Pei et al., 2001). By contrast, hybridization between different clades is less common. For example, seed abortion occurred when *S. exigua* and *S. interior* in clade *Vetrix* were crossed with members of clade *Salix* (Mosseler, 1990). The reproductive barriers could be caused by pollen–pistil incongruity (Mosseler, 1989), and Gulyaev et al. (2022) proposed that different SDSs may also contribute. More recently, genomic approaches were applied to detect hybridization. Sanderson et al. (2023) identified several cases of hybridization between species from different branches among the ancestors of subgenera *Longifoliae*, *Vetrix* and *Chamaetia* (*Vetrix* clade) of *Salix* based on the ABBA-BABA statistics.

According to one version of the Bateson–Dobzhansky–Muller model (BDM model), speciation is expected to happen when reproductive isolation is mostly completed in the absence of gene flow (Hollinger & Hermisson, 2017). However, genetic incompatibilities can be incomplete, allowing some hybridization, or the extent of intrinsic postzygotic reproductive isolation effects may be sensitive to extrinsic environmental circumstances (Cutter, 2023). *Salix* is a case with low genetic divergence between species within the major clades based on restricted DNA barcode markers (Chen et al., 2010), and their relationships within clades can be resolved only with genome-wide data (Gulyaev et al., 2022; Wagner et al., 2020). In such systems where reproductive isolation is still incomplete, the extent of introgression can allow us to uncover whether sex chromosome rearrangements affect the evolution of nascent species boundaries (Wang et al., 2022b). Previous studies of the role of sex chromosome turnovers in genome differentiation have mostly used hybrid swarms created by multi-generation experimental crosses (Volz & Renner, 2008; Wang et al., 2022b). Gene flow studies can be used in a genome-wide phylogenetic framework.

In this study, we sequenced populations of both sexes from *Salix cardiophylla* and *S. interior* and try to **1)** discover their SDSs and determine whether they have XY or ZW system on chromosome 15 or 7; **2)** reconstruct phylogenetic relationships using the SNP data from 15 willow species with known SDSs, and explore whether there are turnover events in the subclades of *Salix* genus and the possible ancestral SDS of *Salix*; **3)** identify willow species with known SDSs that can and cannot undergo gene flow, to get insights whether reproductive isolation has evolved between species that differ in these characteristics.

## Results

### Whole genome re-sequencing

After filtering, we obtained 2,928.19 million clean reads from a natural population of 42 *Salix cardiophylla* individuals of known sex (32.19-37.20 million reads per sample, mean 34.86; Supplementary table 1) and 2,658.38 million reads from 38 *S. interior* individuals (32.10-41.31 million reads per sample, mean 34.98; Supplementary table 2). The average mapping rate varied from 74% to 89.6% (average rate: 85.79%), and the average depth was 29.51× (Supplementary table 3) for *S. cardiophylla*. For *S. interior*, the mapping ratio varied from 89.8% to 92.7% (average rate: 91.53%) and the average depth was 33.71× (Supplementary table 4). We extracted 1,613,125 and 4,060,148 high-quality SNPs from the *S. cardiophylla* and *S. interior* samples, respectively, using *S. purpurea* (chromosome 15 W excluded but 15Z kept, because W degenerated but Z relative conserved) as reference (Zhou et al., 2020).

### Sex determination systems in *S. cardiophylla* and *S. interior*

GWAS and *F*_ST_ results between the male and female samples revealed a sex-linked region on chromosome 15 (15Z of *S. purpurea*) in *S. cardiophylla* and *S. interior* (Fig. 1A, B; Supplementary figure 1, 2, 3, 4), respectively. Changepoint analysis of the *F*_ST_ values detected candidate fully sex-linked regions between 3.59 and 6.42 Mb in the *S. purpurea* 15Z assembly (2.83 Mb, about 21.3% of the chromosome) in *S. cardiophylla*, and 3.32 and 6.35 Mb (3.03 Mb, about 22.8% of the chromosome) in *S. interior* (Fig 1C, D).

**Fig. 1.**
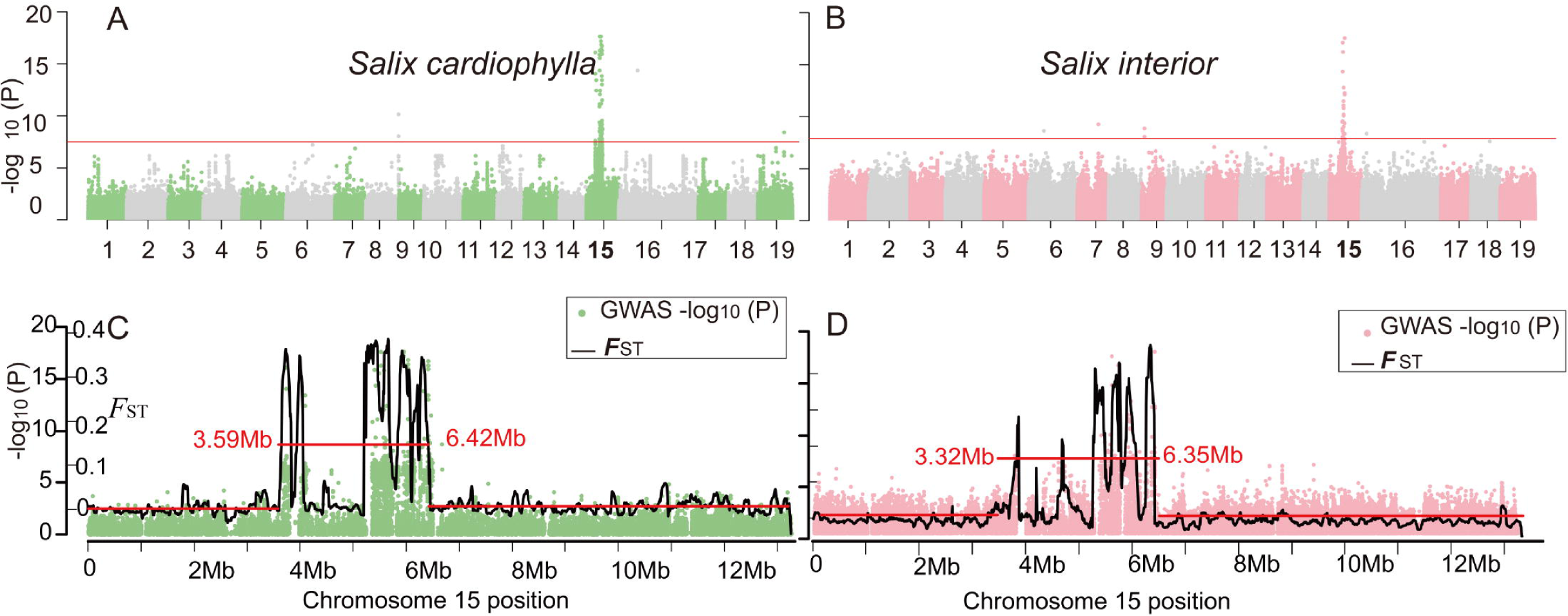
The sex determination systems of *Salix cardiophylla* and *S. interior.* **A**: Genome wide association studies (GWAS) results between SNPs and sexes in 42 individuals for *S. cardiophylla*. **B**: GWAS results between SNPs and sexes in 38 individuals for *S. interior*. The red lines in A and B show the Bonferroni-corrected significance level corresponding to α < 0.05 and the y axis shows the negative logarithm of P values. **C**: P values of GWAS results and *F*_ST_ values between the sexes on chromosome 15 for *S. cardiophylla*. **D**: P values of GWAS results and *F*_ST_ values between the sexes on chromosome 15 for *S. interior*. Red lines represent the significant sex-associated regions analyzed by changepoint analysis both in C and D.

GWAS analysis recovered totals of 78 and 40 SNPs with significant associations with sex on chromosome 15 (15Z of *S. purpurea*) in *S. cardiophylla* and *S. interior*, respectively (Supplementary table 5, 6). In *S. cardiophylla,* 1.29% of these SNPs are heterozygous in females and 33.38% in males. In *S. interior*, 44.61% are heterozygous in the males, and all are homozygous in the females (Table 1). These results suggest that both *S. cardiophylla* and *S. interior* have a sex-linked region on chromosome 15, and male heterogamety.

**Table 1.** Summary of estimated nucleotide homozygosity and heterozygosity rates on sex-linked SNPs based on the GWAS result for *Salix cardiophylla* and *S. interior*.

| Species | Gender | Number | Homozygous(%) | Heterozygous(%) |
| --- | --- | --- | --- | --- |
| *Salix cardiophylla* | Female | 19 | 98.71 | 1.29 |
| *Salix cardiophylla* | Male | 23 | 66.62 | 33.38 |
| *Salix interior* | Female | 20 | 100 | 0 |
| *Salix interior* | Male | 18 | 55.39 | 44.61 |

### Phylogenetic trees based on nuclear and plastid data

Both maximum likelihood (ML) and species tree approaches were applied for the nuclear phylogeny reconstruction: **1)** A phylogenetic tree of 15 willow species with known SDSs and the outgroup poplar species (*Populus euphratica*) was reconstructed using RAxML, based on concatenation of 821,062 high-quality nuclear variants at fourfold degenerate sites. The tree (Fig. 2A) resolved the two main clades, *Salix* and *Vetrix*, with high bootstrap support (100%). The *Salix* clade includes *S. nigra*, *S. chaenomeloides*, and *S. dunnii*. The remaining species fall into the *Vetrix* clade. *Salix cardiophylla* and *S. arbutifolia* formed a subclade. *Salix interior* and *S. exigua* also formed a subclade, which was sister to the other *Vetrix* species, including *S. polyclona*, *S. suchowensis*, *S. koriyanagi*, *S. integra*, *S. purpurea*, *S. viminalis*, and *S. udensis* (Fig. 2A). **2)** For the species tree construction, we reconstructed individual gene trees based on 6,315 variable genes that have at least 50 SNPs. The species tree based on all the genes trees using Astral revealed the same topology as the ML tree (Supplementary figure 5).

**Fig. 2.**
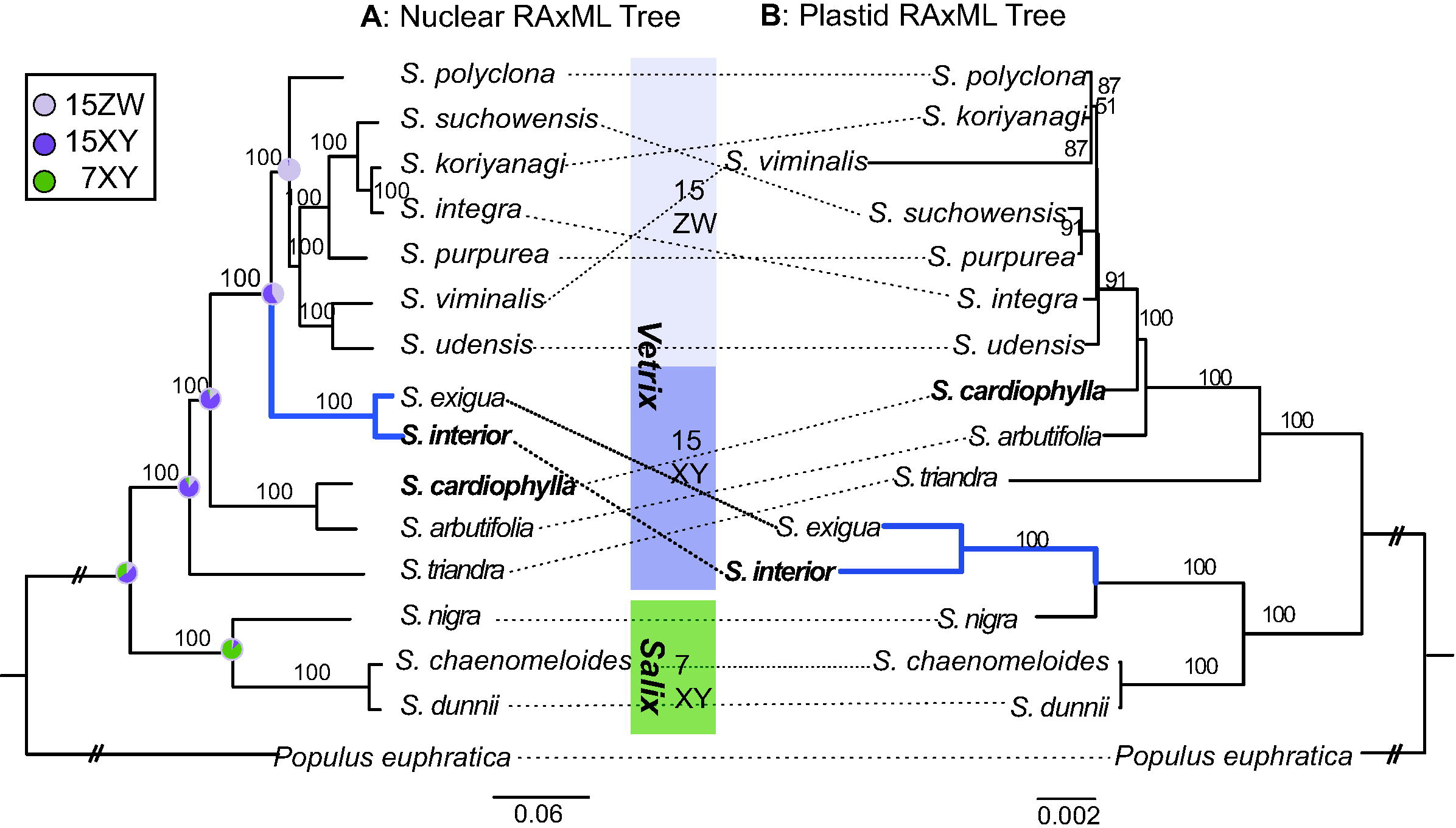
Phylogenetic relationship based on nuclear SNPs (A) and plastid genomes (B) of *S. cardiophylla* and *S. interior* and other *Salix* species with known sex determination systems, using *Populus euphratica* as an outgroup. The numbers at the nodes indicate support values based on 1000 bootstrap replications. The tree is marked with the ancestral character-state reconstruction of sexLdetermining system (light blue: 15ZW; dark blue: 15XY; dark green: 7XY) from Supplementary figure 6.

Trees based on plastid genomes (Fig. 2B) yielded two major well-supported clades, but the subclade including *S. interior* and *S. exigua* belong to the *Salix* clade, rather than the *Vetrix*. *Salix cardiophylla* and *S. arbutifolia* appear as paraphyletic, and basal to a subclade comprising the seven other *Vetrix* species (Fig. 2B).

The ancestral character reconstruction analysis with SDS systems indicated that the most likely ancestral SDS state for the genus *Salix* is 15XY (the relative likelihoods: 53%), while the likelihoods of ancestral 7XY is 35% (Fig. 2A, Supplementary figure 6). For the *Vetrix* clade, with both 15XY and 15ZW SDSs, the most likely ancestral state is 15XY with a proportional likelihood of 81% compared to 15ZW with a proportional likelihood of 11% (Fig. 2A, Supplementary figure 6).

### Gene flow

We applied the heuristic f_branch_ analysis (Malinsky et al., 2018) implemented in Dsuite to measure the level of interspecific gene flow due to hybridization along internal and external branches in *Salix* (Fig. 3; Supplementary table 7). The internal nodes represent hypothetical last common ancestor lineages. The f_branch_ can deliver good accuracy as long as the proportions of simulated gene flow are over 1% (Malinsky et al., 2021). First, the f_branch_ metrics yielded evidence for gene flow between *Salix exigua* and species of both the *Vetrix* clade (f_branch_ = 0.003 to 0.009) and the *Salix* clade (f_branch_ = 0.001 to 0.002). The results also suggested gene flow between the *Salix* clade (with 7XY) and the ancestor of *S. arbutifolia, S. cardiophylla,* and *S. triandra* (f_branch_ = 0.016 to 0.018). Consider the relationship of gene flow between different SDSs systems, 1) the analysis indicated stronger gene flow between species with 7XY and 15XY species (f_branch_ = 0.012 to 0.018) than between 7XY and 15ZW species (f_branch_ = 0.0003 to 0.008, wilcox.test P < 0.01); 2) stronger gene flow were revealed between species with 15XY and 15ZW (f_branch_ = 0.009 to 0.018) than between 7XY and 15ZW species (f_branch_ = 0.0003 to 0.008)(wilcox.test, P < 0.01); 3) Gene flow between species with 15XY and species 15ZW (f_branch_ = 0.009 to 0.018) is a little weaker than the gene flow between 15XY and 7XY species (f_branch_ = 0.012 to 0.018), while the P value is not significant (P = 0.367). Finally, we detected strong gene flow among 15ZW species of the *Vetrix* subclade (0.0009 to 0.064) than the gene flow among either 15XY *Vetrix* clade species (0.006 to 0.009) or 7XY *Salix* clade ones (0.0005 to 0.0006, Supplementary figure 7).

**Fig. 3.**
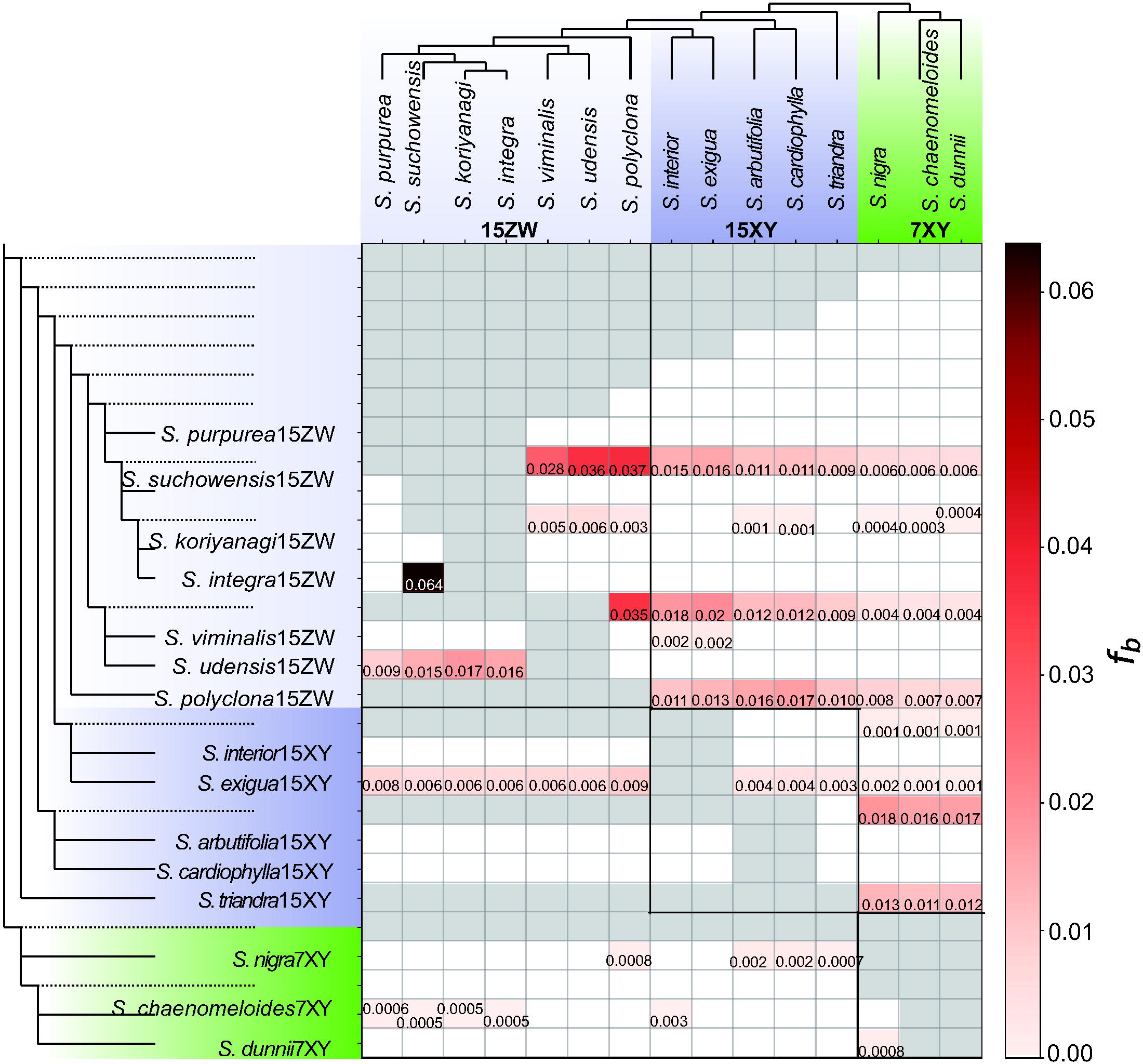
Heuristic f_branch_ analysis results of ongoing and ancient gene flow inferred for species in *Salix*.

## Discussion

Our study confirms that the heterogamety and sex-linked region locations phylogenetically conservative, although changes have occurred between the major groups within *Salix* (Fig. 2). Gene flow rates estimated here are also consistent with previous hybridization experiments (Kimura, 1931, 1937; Mosseler, 1990). This indicates that gene flow detection based on genomic analysis can reliably detect isolation between species, or lack of isolation.

### XY system on chromosome 15 in *Salix cardiophylla* and *S. interior*

Previous studies showed that three of the basal species in the *Vetrix* clade have 15XY systems, including *Salix exigua* (Hu et al., 2022), *S. arbutifolia* and *S. triandra* (Wang et al., 2022a; Wang et al., 2023a). Wang et al. (2023a) suggested that the species of *S. arbutifolia* and *S. triandra* (15XY) are in a transitional stage between 15ZW and 7XY systems. Our results provided two further basal species with 15XY system (Fig. 2).

The American species *S. interior* is considered sister to *S. exigua* and both species have 15XY systems. Both have been classified in section *Longifoliae*, which was considered to be a section of the *Salix* clade (Skvortsov, 1968; Wu et al., 2015). Chen et al. (2010) thought that they belong to the New World subclade of the *Salix* clade. Previous hybridization experiments showed pollen-pistil incongruity of hybrids from *S. exigua*/*interior* and members of the *Salix* clade. By contrast, *S. exigua*/*interior* can hybridize with species of the *Vetrix* clade, and produced fertile *F*_1_ offspring (Mosseler, 1989, 1990). Furthermore, *S. exigua* is among the early-flowering species, similar to many other *Vetrix* clade species, while most willows in the *Salix* clade show late-flowering phenology (Mosseler & Papadopol, 1989). Both phenological and crossing results indicated a closer relationship of *S. exigua*/*interior* with *Vetrix* than with *Salix* clade, which is consistent with Gulyaev et al. (2022) based on phylogenies of nuclear genomic sequences. *Salix interior* grouped with *S. exigua* in our nuclear and plastid genome trees, but the nuclear data place them both in the *Vetrix* clade whereas the plastid sequences assign them to the *Salix* clade in the phylogenetic tree (Fig. 2). This incongruent topology indicates that the two species could have originated from an ancient hybrid between ancestors of the *Salix* and *Vetrix* clades. The phylogeny based on plastid genomes indicates that the maternal lineage may have come from the *Salix* clade, which would imply that a maternal parent from a 7XY species was fertilized by a 15XY/15ZW paternal plant from the *Vetrix* clade. However, this should be further tested.

The species *Salix cardiophylla* was segregated into a separate genus, *Toisusu*, based on its unique morphological characters, including deciduous styles (Kimura, 1928), but molecular phylogenetic analyses assigned it to the genus *Salix* (Chen et al., 2010). Until now, most phylogenetic studies accepted that *S. cardiophylla* is in a clade with *S. arbutifolia* (Chen et al., 2010; Wu et al., 2015). We also revealed *S. cardiophylla* and *S. arbutifolia* form a subclade in the *Vetrix* clade. Wang et al. (2023a) showed that *S. arbutifolia* has male heterogamety and its SDSs is on chromosome 15. Our study confirmed 15XY in *S. cardiophylla*. Although the SDSs are quite changeable in the genus *Salix*, the same SDSs systems of *S. cardiophylla* and *S. arbutifolia* indicated that SDSs within the subclades of *Salix* genus are usually conserved.

### The role of sex chromosome turnovers in speciation of *Salix*

Sex chromosome turnovers play multifaceted roles in the evolution of reproductive barriers among species (Wang et al., 2022b). A fascinating scenario in sex chromosome evolution connected to the process of speciation is hybridization, especially hybridization of species with different heterogamous sex chromosomes. Limited introgression was observed in species of different sex determination systems (Baack et al., 2015). Franchini et al. (2018) conducted a long-term evolutionary crossing experiment of two fish species with ZW and XY SDSs, and demonstrated that reproductive isolation present but incomplete between different SDSs. Kitano and Peichel (2012) demonstrated the reproductive isolation between multiple sex chromosome systems and proved the important role of sex chromosome turnover in the speciation of sympatric stickleback species. Similar patterns were found in plants, the neo-sex chromosomes serving as introgression barriers between diverging species of *Rumex* (Beaudry et al., 2020). Based on current datasets, we revealed turnovers in *Salix* genus, i.e., 15XY➔7XY and 15XY➔15ZW, consistent with the finding in (Wang et al., 2023b). Our *Salix* gene flow results further suggested that different SDS systems in *Salix* genus possibly act as the barrier to introgression in varying degrees (Fig. 3). However, 15XY to 15ZW and 15XY to 7XY, that carried same sex determination factors(Wang et al., 2023b), are insufficient to form a complete reproductive barrier between the species pair, at least in early generations of their separation. However, Sanderson et al. (2023) found a burst of diversification near the origin of the *Vetrix* clade, which is likely match the SDSs 15XY➔15ZW transition in this clade.

Our analysis found that little gene flow occurs between the species having 7XY and 15ZW SDS, suggesting that the trans-heterogamety transitions might play an important role in willow speciation, which is also found in stickleback fish (Kitano et al., 2009). However, we cannot rule out that incompatibilities between the *Salix* clade and the *Vetrix* clade are due to overall genetic divergence, which is much larger between the big clades than within them (Fig. 2). It was suggested that neo-sex chromosomes from the turnover events contributed to ecological specialization and speciation process in *Drosophila* (Yu et al., 1997). Although turnover of sex chromosomes of willows may not act as a complete reproductive barrier (Gulyaev et al., 2022; Stock et al., 2021), the different SDSs could contribute to reproductive isolation and facilitate the speciation process (Olito & Abbott, 2023). In our study, historic introgressions were observed between the ancestor of 15ZW and 15XY species, while contemporary gene flow between the species with the two SDSs is obviously reduced (Fig. 3). The present genomic data suggests that after the turnover, SLRs in *Salix* genus have evolved independently in different lineages during the subsequent evolutionary process as revealed by Wang et al. (2023b). Studies revealed that incompatible loci/micro-heteromorphism (see below) for hybrid male/female sterility likely accumulated on the sex-linked regions on the sex chromosomes, which likely further facilitate the reproductive isolation in different subclade of *Salix* and thus promote the speciation (He et al., 2023b; Wang et al., 2023b; Zhou et al., 2020).

Sex chromosomes pairs in plants are thought to be homomorphic and young (Wang et al., 2022a). Homomorphic sex chromosomes do not differ in size at the karyotype level and usually have a relatively small non-recombining region. However, recent studies showed a high diversity in the size of sex-determining regions across plants, from a single locus in poplar (Müller et al., 2020) to a small differentiated region in strawberry (Tennessen et al., 2018) and heteromorphic sex chromosomes with large non-recombinant regions in *Silene* (e.g., Filatov, 2022). Changes in the size of sex-linked regions can shed light on how sex chromosomes evolved, including how their suppression of recombination arose, as well as the timeline of subsequent evolution (Long et al., 2023). The SLRs varied among different species of *Salix*, which may relate to the sex chromosome systems. Willow species with known sex-linked region lengths, using *S. purpurea* as a reference: *Salix triandra* (15XY) SLR 2.8 Mb (Wang et al., 2023a), *S. arbutifolia* (15XY) SLR 3.33 Mb, *S. cardiophylla* (15XY) SLR 2.83 Mb (this study), *S. interior* (15XY) SLR 3.03 Mb (this study), *S. purpurea* (15ZW) Z-SLR 4.4 Mb (Zhou et al., 2020), and *S. polyclona* (15ZW) SLR 4.68 Mb (He et al., 2023a). These high variations of the sex-linked region in *Salix* may also indicate the independent evolution in the sex determination systems, like in the true frogs (Jeffries et al., 2018), and sticklebacks (Yoshida et al., 2014). The divergence time of *S. cardiophylla* and *S. interior* with 15XY SDS is earlier than that of other species in the *Vetrix* clade with 15ZW SDS (Wu et al., 2015), but by comparing the size of the SDR, the sex-linked regions of *S. cardiophylla* and *S. interior* were smaller although they have had a longer time for SDS evolution, which further indicate SDSs evolution in *Salix* is independent.

Furthermore, with available phased genomes, micro-heteromorphism between X/W and Y/Z was detected in three species distributed in different clades covered all kinds of willows’ SDSs (table 2): *S. dunnii* 7XY (He et al., 2023b), *S. arbutifolia* 15XY (Wang et al., 2023b), *S. purpurea* 15ZW (Zhou et al., 2020). In detail, 7X-SLR of *S. dunnii*, 15X-SLR of *S. arbutifolia*, and 15W-SLR of *S. purpurea* accumulated more repeat sequences comparing to their homologous X- or W-SLR, and different specific genes were identified on the X-Y and Z-W pairs (Wang et al., 2023b). During the diverge process from initially identical sex chromosomes to heteromorphic sex chromosomes, sequence loss, the accumulation of deleterious mutations sex bias genes and repetitive elements might reduce the gene flow in sex chromosome and thus play the reproductive factors during the speciation (Ping et al., 2022). The heteromorphic sex chromosomes play important roles in hybrid sterility/inviability (Drosophila, Ferree & Barbash, 2009) and in reducing introgression of sex chromosome compared with the autosomes in *Heliconius* butterflies (Martin et al., 2013) and *Ficedula* flycatchers (Ellegren et al., 2012), while the homomorphic fish species didn’t show X-autosome introgression differences (Schumer et al., 2014). In Salicaceae, the sister genus *Populus* with the small size of the SDS in species with known SDSs (0.08 to 2 Mb, Wang et al., 2023a) and a single-locus system with epigenetic control (Müller et al., 2020), which probably facilitate hybridizations between species with ZW and XY systems, e.g., *P. alba* ZW x *P. tremula* XY = *P. x canescens*, an abundant hybrid species which is fully fertile (Meikle, 1984; Zhang et al., 2023). In *Salix*, the hybridization between different clades (*Salix* and *Vetrix*) of *Salix* is less common (Mosseler, 1990), which may contribute from the long sex-linked regions and the micro-heteromorphic chromosomes since sex-linked regions can accumulate more non-random sex specific or the incompatible recessive genes. These will further reduce the recombination rate and hybrids are more likely to suffer from infertility. However, to which extent the SDSs in *Salix* inhibits reproductive isolation and promotes the speciation is comparable needs to be further studied.

**Table 2.** *Salix* Species with phased chromosomes.

| Species | SDS systems | X/W length (Mb) | SLR length (Mb) | SLR Percent | Y/Z length (Mb) | SLR length (Mb) | SLR Percent | References |
| --- | --- | --- | --- | --- | --- | --- | --- | --- |
| *S. dunnii* | 7XX/X<br>Y | 19.21 | 5.87 | 30.56% | 16.39 | 2.95 | 18.00% | He et al., 2023b |
| *S. arbutifolia* | 15XX/<br>XY | 16.16 | 7.06 | 43.69% | 10.94 | 1.81 | 16.54% | Wang et al., 2023b |
| *S. purpurea* | 15ZZ/Z<br>W | 15.65 | 6.7 | 42.81% | 13.3 | 4.4 | 33.08% | Zhou et al. 2020 |

## Methods

### Taxa sampling

For the whole-genome resequencing, we collected 42 individuals for diploid *Salix cardiophylla*, including 19 female and 23 male individuals from Japan (Supplementary table 8). For diploid *S. interior,* we sampled 38 individuals, including 20 female and 18 male individuals from America (Supplementary table 8). The two species were identified based on the descriptions in relevant floras and taxonomic studies (Argus, 2010; Ohashi, 2006). For each individual, leaves were dried with silica gel and the voucher specimens were deposited at the herbarium of Shanghai Chenshan Botanical Garden (CSH).

To confirm the systematic position of the two species in our study, we downloaded whole genome sequence data of 13 representative species, with known SDSs, of *Salix* genus (Supplementary table 9), which have available genome data. The species *Populus euphratica* (SRR13324572) was also included as an outgroup in this study.

### Sequencing, reads mapping, and variant calling

Total genomic DNA of *Salix cardiophylla* and *S. interior* individuals were isolated from silica-dried leaf samples using Qiagen DNeasy Plant Mini Kit (Qiagen, Valencia, CA) following the manufactures’ instructions. The whole-genome paired-end sequencing library was constructed and sequenced on Illumina NovaSeq 6000 by Beijing Novogene Bioinformatics Technology, China.

The raw sequenced reads were filtered by using fastp 0.20.0 (Chen et al., 2018). Clean reads of *Salix cardiophylla* and *S. interior* were aligned to the closed related published reference genome of *Salix purpurea* (Zhou et al., 2020, chromosome 15 W excluded but 15Z kept) by BWA 0.7.12 with default settings (Li & Durbin, 2009). We then extracted primary alignments, sorted, and merged the mapped files in SAMtools 0.1.19 (Li et al., 2009). Sambamba 0.7.1 (Tarasov et al., 2015) was performed to discard clonal duplicates during the library preparation.

We called the variants using the program ‘HaplotypeCaller’ and ‘GenotypeGVCFs’ in Genome Analysis Toolkit (GATK) v. 4.1.8.1 (McKenna et al., 2010). In detail, we used the setting ‘–sample-ploidy 2’ in ‘HaplotypeCaller’ for the chromosome regions.

Hard filtering was used for the further SNP calling, with the setting (QD < 2.0, FS > 60.0, MQ <40.0, MQRankSum < −12.5, ReadPosRankSum < −8.0, SOR > 3.0). We only kept the biallelic sites in the chromosome regions. For the subsequent filtering, sites with coverage more than twice the average depth at variant sites in all samples were excluded. Genotypes with depth <4× were treated as no-call, sites with no-call genotypes in more than 10% of samples were removed, and sites with minor allele frequency < 0.05 were further discarded. We also used the same method to call SNPs for phylogenetic analysis.

### Identification of the sex determination system of *S. cardiophylla* and *S. interior*

To identify the sex determination system of the two species, we first extracted high-quality SNPs, from 42 *S. cardiophylla* and 38 *S. interior* individuals, respectively. The two datasets were performed in a standard case-control genome wide association studies (GWAS) of allele frequencies and sex phenotypes with PLINK v1.90b6.18 (Chang et al., 2015). SNPs with α < 0.05 after Bonferroni correction for multiple testing yielded 78 and 40 sex linked-SNPs on chromosome 15Z, respectively, in *S. cardiophylla* and *S. interior*.

We also calculated the genetic differentiation (*F*_ST_) between the male and female individuals in VCFtools 0.1.16 (Danecek et al., 2011) using the Weir and Cockerham (1984) estimator with 100-kb windows and 10-kb steps. Regions with significantly higher *F*_ST_ values than other parts of the chromosome are considered candidate sex-linked regions (He et al., 2021). A Changepoint package (Killick & Eckley, 2014) was used to assess significance of differences in the mean and variance of the *F*_ST_ values in chromosome 15 windows, using the function cpt.meanvar, algorithm PELT and penalty CROPS.

VCFtools was then used to calculate heterozygote frequencies of sex linked-SNPs detected by GWAS on a per-individual basis. High heterozygosity in males compared with females suggests male heterogamety, and higher heterozygosity in females indicates female heterogamety.

### Plastid genome assembly and alignment

We assembled 16 plastid genomes using Getorganelle v 1.7.6.1 (Jin et al., 2020) from the clean sequence data with default parameters. Homblocks v 1.0 (Bi et al., 2018) was then employed to align the sequences for the following phylogenetic analysis.

### Phylogenetic analysis and ancestral SDS reconstruction

To infer the phylogenetic relationship of *S. cardiophylla* and *S. interior* in the *Salix* genus, we included the genomic sequences of another 13 representative *Salix* taxa and one species of *Populus* as outgroup. We used the genome of *S. purpurea* as the reference again and called the single nucleotide variants (SNPs) after excluding the putative sex chromosomes 7, and 15. This yielded out 10,665,405 SNPs. The SNPs that are fourfold degenerate sites in the reference genome were detected by a python script based on the gene annotation of *S. purpurea* (https://github.com/zhangrengang/degeneracy), and only 821,062 fourfold degenerate sites SNPs were kept. Finally, phylogenomic relationships based on the chromosome dataset were reconstructed by a maximum likelihood approach using RAxML v.8.2.4 (Stamatakis, 2014). Support values were calculated using 1000 rapid bootstrap replicates based on the GTR + GAMMA nucleotide substitution model. We applied the same settings as the chromosome dataset to infer the plastid RAxML tree.

To infer the relationship among the 16 species, we also estimated species trees. We used Orthofinder 2.5.2 (Emms & Kelly, 2019) to identify single-copy genes in all five diploid genome assemblies of *S. brachista* (Chen et al., 2019), *S. dunnii* (He et al., 2021), *S. purpurea* (Zhou et al., 2020), *S. suchowensis* (Dai et al., 2014) and *S. viminalis* (Almeida et al., 2020). We chose the more variable genes with at least 50 SNPs in the coding regions of each gene from the 10,665,405 SNPs dataset. Modelfinder (Kalyaanamoorthy et al., 2017) was then used to select the best model based on Bayesian information criterion, and IQ-tree v. 2.1.4 (Minh et al., 2020) was applied to reconstruct individual gene trees based on the selected best model and ascertainment bias correction model. A species tree was estimated using Astral v. 5.7.8 (Zhang et al., 2018), and clade support was calculated using local posterior probabilities (Sayyari & Mirarab, 2016). The species tree was then used for Dsuite analysis.

To test the possible ancestral states of SDSs in willows, we reconstructed the characters ancestral states in Mesquite v.2.0 (Maddison, 2007) based on the RAxML tree from the chromosome dataset. In detailed, the 7XY systems were coded as 0, 15XY systems were coded as 1, and 15ZW systems as 2 in the probability model.

### Gene flow in the genus *Salix*

To test the gene flow across the related species in *Salix*, we calculated ABBA-BABA (Patterson’s D), f_4_ and f_branch_ statistic in Dsuite 0.5 package (Malinsky et al., 2021). The statistic of f_4_-ratio provides a format test for the admixture history. The f_branch_ statistic is heuristic and is applied to account for the phylogenetic correction among f_4_ ratio results and assign the gene flow to the external (current) and also the possibly internal branch (the hypothetical ancestor lineages). The VCF file (1066,5405 SNPs) generated by GATK and the species tree (see above) were used as input files for Dsuite analysis. The significance of each test was assessed using 100 jackknife resampling runs.

## Acknowledgements

This study was financially supported by the National Natural Science Foundation of China (grant no.32171813), Special Fund for Scientific Research of Shanghai Landscaping & City Appearance Administrative Bureau (grant nos. G232403 & G242417), and National Wild Plant Germplasm Resource Center for Shanghai Chenshan Botanical Garden (ZWGX2202). We are grateful to Satoshi Kikuchi for sampling, to Natalie Konig for assistance with field work, and to the Chesterfield, Missouri Parks, Recreation and Arts Department for permission to collect. We thank Deborah Charlesworth, Yi Wang, Yuàn Wang, Hui Shang and Guangnan Gong for the helpful comments.

## Author Contributions

L.H. designed the study; L.H and Z.X. performed the analysis and wrote the manuscript with input from W.L.A. and E.H.; All authors contributed to later versions of the manuscript.

Conflict of interest: The authors declare no conflict of interest.

## Data availability statement

Sequence data presented in this article can be downloaded from the NCBI database under BioProject accession PRJNA984197 for *Salix cardiophylla* and PRJNA984212 for *Salix interior*.

## Supplementary

**Supplementary table 1**. Summary of sequencing quality of samples of *Salix cardiophylla*

**Supplementary table 2**. Summary of sequencing quality of samples of *Salix interior*

**Supplementary table 3**. The mapped results of the high-quality reads of all 42 samples of *Salix cardiophylla* using *Salix purpurea* as the reference genome

**Supplementary table 4**. The mapped results of the high-quality reads of all 38 samples of *Salix interior* using *Salix purpurea* as the reference genome

**Supplementary table 5**. The 78 sex-linked SNPs on chromosome 15 of *Salix cardiophylla* using *Salix purpurea* as the reference genome

**Supplementary table 6**. The 40 sex-linked SNPs on chromosome 15 of *Salix interior* using *Salix purpurea* as the reference genome

**Supplementary table 7.** F_branch_ metrics in *Salix* using the ASTRAL species tree as input tree

**Supplementary table 8** Details of the collection information of 42 samples of *Salix cardiophylla* and 38 samples of *S. interior*

**Supplementary table 9**. The species of willows with known sex determination system in this study

**Supplementary figure 1.** Quantile–Quantile (Q–Q) plots of observed and expected GWAS *P*-values in *Salix cardiophylla*. Red dotted line indicates X = Y and blue shading the 95% confidence interval around the expectation of X = Y

**Supplementary figure 2.** Quantile–Quantile (Q–Q) plots of observed and expected GWAS *P*-values in *Salix interior*. Red dotted line indicates X = Y and blue shading the 95% confidence interval around the expectation of X = Y

**Supplementary figure 3.** The genetic differentiation (*F*_ST_) between the male and female individuals of *Salix cardiophylla*

**Supplementary figure 4.** The genetic differentiation (*F*_ST_) between the male and female individuals of *Salix interior*

**Supplementary figure 5.** Inferred species tree of 15 willow species and the outgroup *Populus euphratica* in this study

**Supplementary figure 6.** The reconstruction of ancestral sex determination system of willow species with known SDSs information

**Supplementary figure 7.** The comparison of F_branch_ metrics within the willows with same sex determination system (A) and across willows with different sex determination systems (B) genus with known sex determination systems using Dsuite based on the SNP dataset. The nuclear species tree is shown at the top of the matrix. The tree is displayed in an “expanded” form along the y axis points to a corresponding row in the matrix with ancestral branches. The values in the matrix refer to excess allele sharing between the expanded tree on the y-axis and the species tree on the x-axis.

## Notes

### Competing Interest Statement

The authors have declared no competing interest.

